# The integration of tactile and proprioceptive signals to achieve haptic object perception

**DOI:** 10.1101/2023.11.27.568836

**Authors:** R. Efe Dogruoz, Natalya S. Shelchkova, Drew E. Sheets, Charles M. Greenspon, Sliman J. Bensmaia

**Affiliations:** Department of Psychology, Yale University, New Haven, CT; Department of Organismal Biology and Anatomy, University of Chicago, Chicago, IL

## Abstract

Stereognosis, the sense of the 3-dimensional shape of objects held in hand, requires the integration of somatosensory signals about local features -such as edges and surface curvature- with proprioceptive signals about the conformation of the fingers on the object. However, the mechanism of this integration remains unknown. Here, we investigated the spatial model that is used to integrate information about the global shape of the object with information about its local features at each point of contact. To this end, human observers judged the dissimilarity of pairs of objects that differed in their global shape, their local features, or both. We then compared the dissimilarity ratings when both global shape and local features changed to ratings when only global shape or only local features changed. We tested this with object sets of different levels of complexity, including spheres of different sizes and surface features to more varied shapes and features. For all object sets, we found that ratings when both global shape and local features changed was approximately an additive combination of the ratings when only global shape or only local features changed. For the majority of subjects, a city-block spatial model best explained their responses. Our results suggest that information about global shape is encoded independently from that about local features during interactions with objects. This implies that the neural representations of object shape and local features, though integrated, are separable.

## Introduction

During interactions with objects, cutaneous mechanoreceptors detect skin deformations and transduce them into neural signals that are then transmitted to the brain. The mapping between skin deformations and these signals is well-characterized, allowing us to decode texture and contact force from the population activity of tactile afferents (Saal et al., 2017, 2018; Graczyk et al., 2016). In contrast to these single-modality sensations, stereognosis - the sense of the 3-dimensional shape of objects held in hand - requires the integration of signals from two modalities: tactile signals about local features at each point of contact (such as edges or curvature); and proprioceptive signals about the locations of contact points in space, without which the tactile signals cannot be interpreted (Delhaye et al., 2018). Tactile and proprioceptive signals are relayed to the brain by different peripheral nerves, and how they are integrated to give rise to a shape percept is unknown. Here, we attempt to shed light on this integration in a psychophysical experiment.

Information about touch -such as force, vibration or scanning speed-is conveyed by cutaneous nerve fibers that innervate the skin (SA1, SA2, RA, and PC receptors; Darian-Smith, 1984). Afferent signals project to the cuneate nucleus in the brainstem, then to thalamus, from where they are relayed to Brodmann’s area 3b of somatosensory cortex (S1; Delhaye et al., 2018). Proprioceptive signals are encoded by muscle spindles and Golgi tendon organs in muscle fibers, and ultimately project to area 3a of S1 via cuneate nucleus and thalamus. Tactile and proprioceptive streams are not integrated until area 2 of S1. This integration is thought to be necessary for stereognosis, but the phenomenon has received little experimental attention to date, and mechanisms underlying it remain unknown (Norman et al., 2004; Lacey et al., 2007).

To fill this gap, we had human observers judge the dissimilarity of object pairs that differed in global shape (proprioceptive inputs), local features (tactile inputs), or both at the same time. We asked how the perceptual dissimilarity driven by differences in shape was combined with that driven by differences in features to drive the dissimilarity when both dimensions changed simultaneously.

One way to test multimodal integration is by testing spatial models that describes the relationship between bimodal and unimodal distances (Garner & Felfoldy, 1970; Hyman & Well, 1967). Past work has argued that “separable” dimensions, dimensions which either one can be attended to while ignoring changes in the other, yield City-Block distances where bimodal distances are an arithmetic sum of unimodal ones, while integral dimensions yield Euclidean distances which rely on the Pythagorean theorem (Garner & Felfoldy, 1970). For example, in vision, it has been shown that color and shape are separable, while value and chroma are integral dimensions (Hyman & Well, 1967). Therefore, we sought to answer which spatial model (e.g., City-Block or Euclidean) best describe the integration of tactile and proprioceptive signals during stereognosis.

We used a dissimilarity task over alternatives like object recognition or haptic search (van Polanen et al., 2014) for two reasons: First, we reasoned that dissimilarity tasks would be less susceptible to pop-out effects where specific features of objects can be used as cues to make judgments, thus ensuring that subjects attend to the whole object. It has been shown that pop-out effects indeed exist for both alternatives (Norman et al., 2019; van Polanen et al., 2014). Second, the dissimilarity rating paradigm can be used to quantitatively assess how two sources of information are combined, unlike recognition rates or search time, which exhibit a much more complex relationship with the sensory evidence.

We investigated the integration of tactile and proprioceptive signals using three sets of objects with different levels of complexity. Across all three object sets, we found that dissimilarity when both local features and global shape change can be predicted from a simple additive combination of the ratings when only local features or only global shape changes. Further, for most participants, the best-fitting spatial model was closer to City-Block than Euclidean. These results held true even when the object space was too complex for conscious memorization of ratings. We conclude that local features and global shape are represented surprisingly independently during haptic object perception.

## Results

To investigate the integration of tactile and proprioceptive contributions to stereognosis, we had human subjects perform a dissimilarity task. Briefly, subjects were seated at an experimental table where they could not see the objects (Figure 1). On each trial, subjects were presented with a pair of objects, which they grasped in succession, and then report the dissimilarity between the objects. Dissimilarity was defined as how different the pair of objects were *overall*, with identical objects ascribed a rating of zero. Subjects grasped the object with their thumb, index, and middle fingertips by flexing their fingers (Figure 1d) and without moving their fingers around the object. Each object was grasped for approximately three seconds, and the time between the presentation of two paired objects was approximately 2-4 seconds.

**Figure 1.**
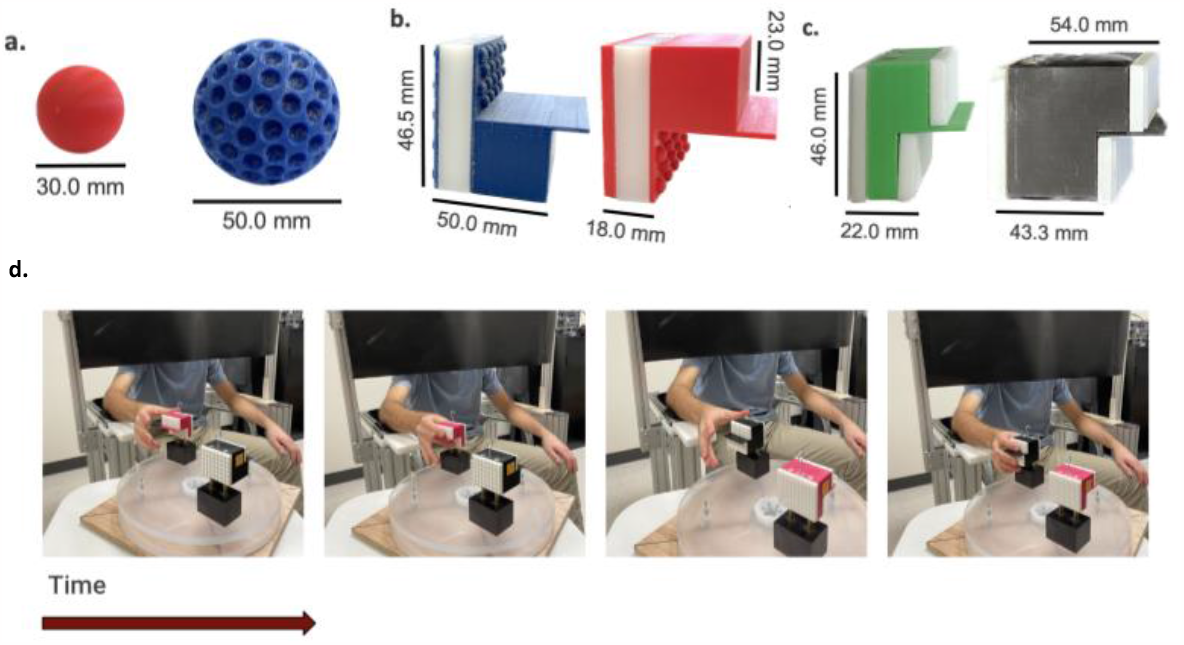
Dissimilarity task. a - c | Object samples used in C1, C2, and C3, respectively. d | Trial progression. In each trial, the subject grasped the first object for ∼3 seconds, released after cue, grasped the second object for ∼3 seconds, and gave a rating of dissimilarity.

We tested how local features and global shape are integrated using three sets of objects with different levels of complexity. C1 objects were plastic spheres of three sizes (diameters 3 - 5 cm), whose surfaces were tiled with one of three local features (bumps, holes, or smooth surface; see Methods; Figure 1a; Figure S1). C2 objects were rectangular blocks (5.1 × 4.65 × 1 or 4 cm) that had one of three global shapes (“L” shape, inverted-”L” shape, and “I” shape). The subjects grasped these objects such that the thumb would contact one side and the index and middle fingers would contact two surfaces on the opposite side of the object. Different surfaces on the same object could have different local features (which were either bumps or smooth surfaces). C3 objects were similar in shape to their C2 counterparts but were more varied to preclude memorization of the object pairs and their associated rating. The objects were rectangular blocks with one of eight global shapes (with different degrees of thumb-index and thumb-middle finger separations), and one of six types of local features, which were distributed uniformly on the contact surfaces.

A total of 19 subjects participated in the study (five subjects in C1, six subjects in C2, and eight subjects in C3). Pairs were presented in randomized order and each pair was presented 3-5 times across different blocks, depending on the object set. On some trials, the same object was presented twice to obtain a baseline level of dissimilarity. Because the subjects chose their own scales, raw dissimilarity ratings were normalized to the mean dissimilarity rating of the corresponding block. For each pair, a dissimilarity score was obtained by averaging the normalized dissimilarity ratings across blocks.

### Subjects Achieve High Signal-to-Noise

First, we assessed whether subjects were following task instructions and able to perform the task by comparing the mean rating across four types of pairs: Pairs in which only features changed, pairs in which only shape changed, pairs in which both changed, and pairs in which neither changed (so called ‘Same’ pairs). We found that ‘Same’ pairs were ascribed lower dissimilarity scores than ‘Feature’, ‘Shape’, or ‘Both’ pairs, consistent with expectation (Figure 2). This held true for all 17 subjects that were presented with Same pairs.

**Figure 2.**
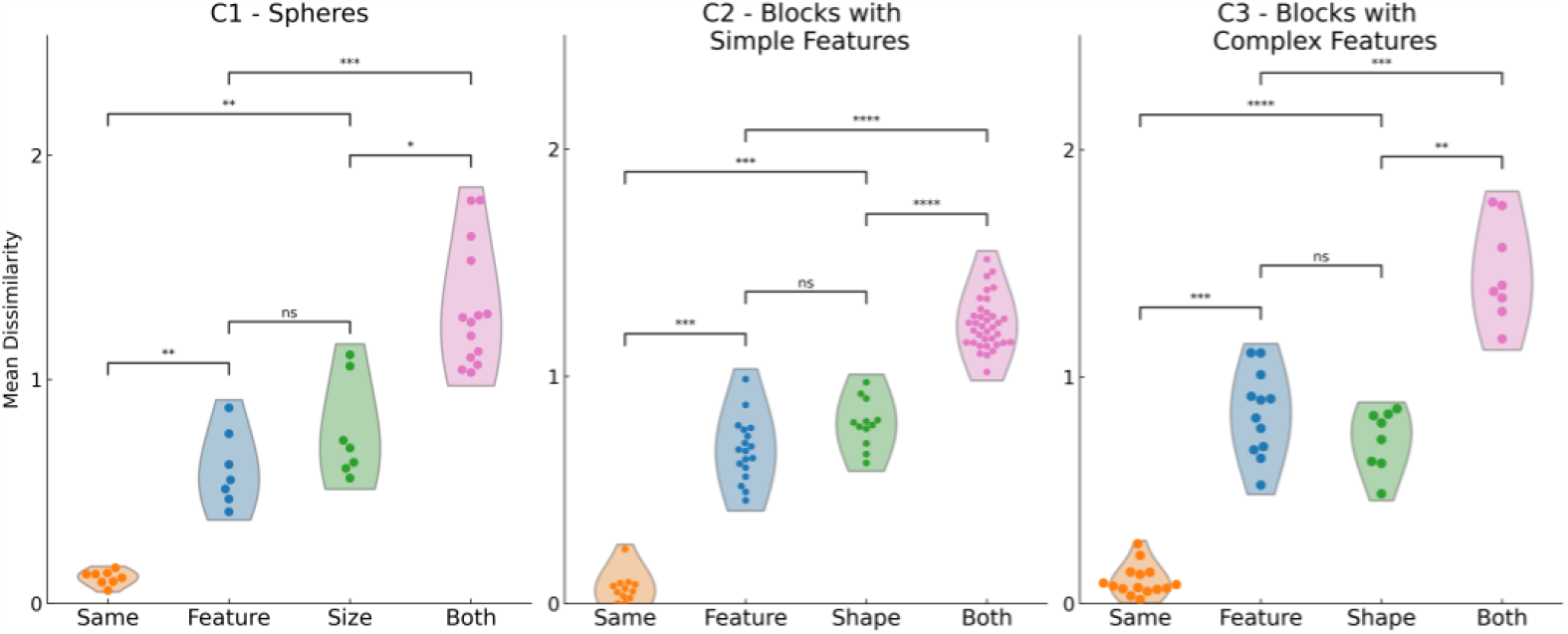
Contribution of global shape and local features to perceived dissimilarity. For all object sets, subjects correctly rated the Same pairs as being the least dissimilar. Changes in shape and local features had a comparable impact on dissimilarity for all object sets. When both local features and global shape changed, subjects gave higher dissimilarity scores on average compared to when only shape or only local features changed (p < .05). Ratings are compared with a two-sided Mann-Whitney-Wilcoxon test and Bonferroni corrected for multiple comparisons. * = p < .05, ** = p < .01, *** = p < .001, **** = p < .0001.

Second, because each different pair entailed a unique change, we next asked if the subjects rated some pairs as being more dissimilar than others. For this, we computed a signal-to-noise ratio (SNR) metric by dividing the variation in dissimilarity scores across different pairs by the average variation in dissimilarity ratings within pairs (see Methods). This metric gauged whether a subject’s ratings varied more between different pairs than between repeated presentations of the same pair. As expected, the SNR metric was negatively correlated with another measure of noise, namely the deviation from zero of the ratings ascribed to the Same pairs (Figure S2). To the extent that ratings varied across different pairs more than within each pair, the SNR was greater than one. The SNRs were greater than 1 for all but one of the subjects. This subject also had the highest average dissimilarity rating for Same pairs, and was thus excluded from further analysis (Figure S2).

Next, we examined to what extent the perceived dissimilarities were consistent across subjects. This analysis yielded average coefficient correlations of 0.68, 0.38, and 0.31, for C1-C3, respectively, highlighting how the increasing complexity of the object sets results in more subjective perceptions of the presented pairs (Figure S3). In conclusion, although subjects were consistent in their ratings, they did not always agree with one another. Thus, we did not pool subjects for further analyses.

### Changes in Local Features and Global Shape are Integrated Additively

We first assessed if the average dissimilarity ratings differ by pair type (Figure 2). As expected, Same pairs yielded the smallest dissimilarity scores for all object sets. Changes in local features alone or global shape alone had similar impacts on perceived dissimilarity on average (difference not significant for C1-3), though this varied across individual subjects, with some rating Feature higher than Shape pairs and vice versa. Finally, pairs where both features and shape changed were perceived to be more dissimilar than pairs where only features or only shape changed for all object sets (p < .05, Mann-Whitney-Wilcoxon test). This was true for all 18 subjects (p < .05, Mann-Whitney-Wilcoxon test). Thus, in line with our expectations, when both local features and global shape changed simultaneously, subjects gave higher ratings of dissimilarity than when only local features or only global shape changed.

We then asked how the specific feature and shape changes contributed to dissimilarity when both dimensions changed simultaneously. If both dimensions are weighted equally and combined additively, the sum of the dissimilarity ratings given to Feature-only and Shape-only pairs should equal the rating given to the corresponding Both pair. To test this, we ran a linear regression model predicting the Both pair ratings from the sums of the corresponding Feature and Shape pair ratings, separately for each subject. To the extent that Feature and Shape changes were combined additively, the coefficient of the summed values would be close to 1. Indeed, the coefficient values were centered close to 1 for all object sets, with median values of 0.99, 0.94, and 0.92 for C1-3 (Figure 3a), showing that the additive model was able to predict the Both ratings to a reasonable degree. Consistent with previous literature (Hyman & Well, 1967), the coefficients varied across subjects, with some being subadditive and others superadditive, although they were centered around the additive coefficient of 1.

**Figure 3.**
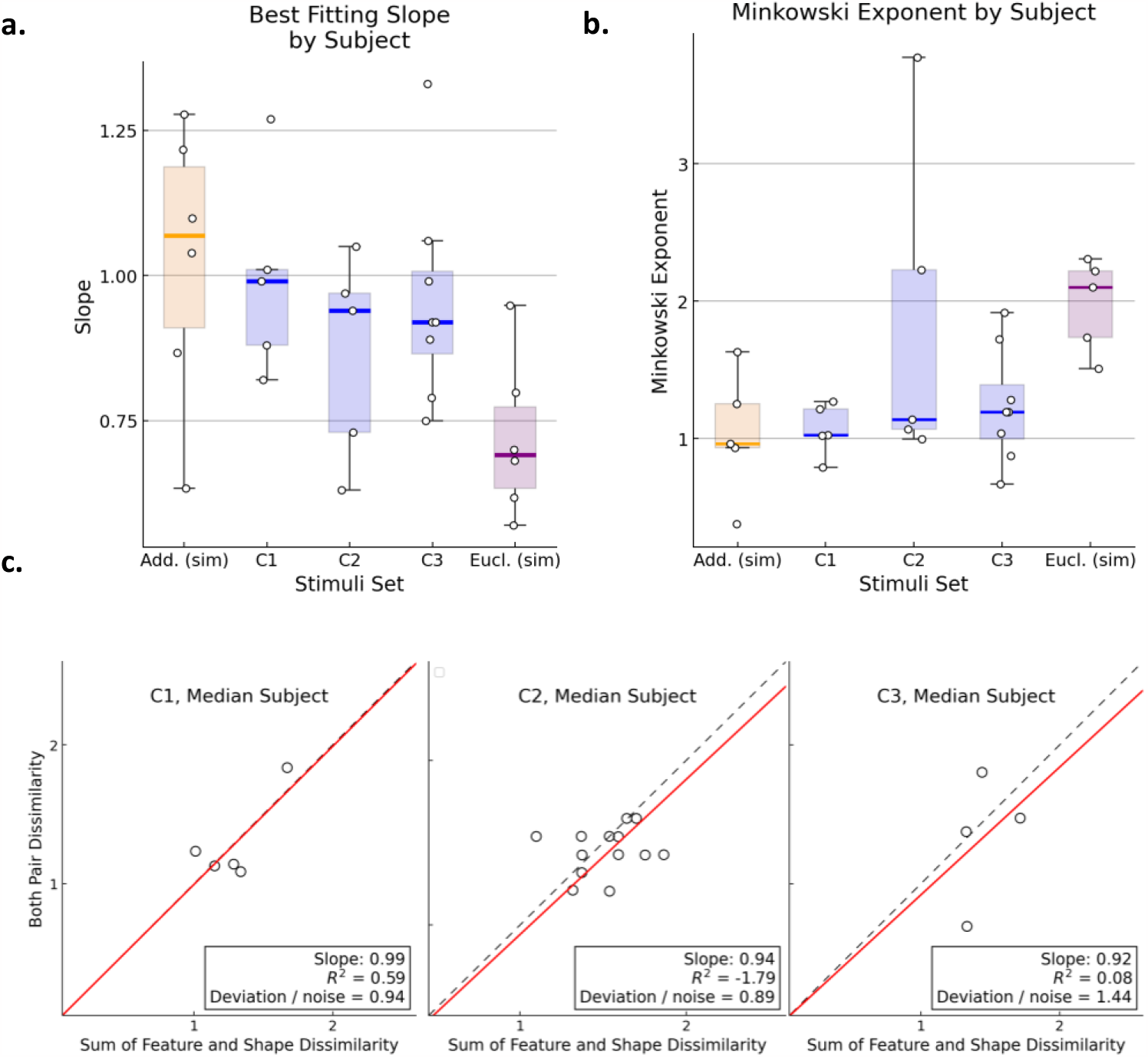
Global shape and local features are integrated additively and in a City-Block manner. a | Slope of the regression model predicting Both pair ratings from a sum of the corresponding Shape and Feature ratings by subject and stimuli set. For both a and b, simulated data for an additive model (orange) and Euclidean model (purple) are shown for comparison. b | Best-fitting Minkowski exponent for each subject and stimuli set. Subjects center close to an exponent of 1 indicating a City-Block spatial model. c | Slopes show the weight of the summed values for predicting the Both pair ratings in a linear regression model (red line) through the origin. Deviation refers to the RMSE of the additive model (dashed lines), and noise is the average root mean squared deviation from the mean rating for each pair.

Because the coefficient of determination depends on the spread of the data and not only on the match, we also tested goodness of fit with a second performance metric, namely the ratio of the deviation from the additive model (RMSE) to the intrinsic noise, defined as the mean RMS deviation from the mean for all different pairs (see Methods). RMS deviation was computed separately for each subject, then averaged across subjects. This metric (“Deviation / noise”) yielded values of 0.94, 0.89, and 1.38 for the subjects with median coefficient values in object sets C1-3, suggesting that the deviation from additivity was comparable to or smaller than the noise in the data for these subjects (Figure 3c). These results are thus consistent with the hypothesis that the Both ratings are approximately the sum of their associated Feature-only and Shape-only ratings.

As the Deviation / noise metric produced a range of values above and below 1 across subjects, we next assessed which of these values were significantly different from what is expected from an additive model. For each subject, we performed a Monte Carlo simulation where we created a distribution of possible error values under the assumption of perfect additivity given the amount of noise in the data (Figure S4; see Methods). The measured error did not differ from the error values simulated assuming additivity for more than half (10/18) of the subjects (Figure S4). All but two of the eight subjects who deviated from additivity could be explained with a linear model where the sum of Feature and Shape ratings were factored by a coefficient, either sub-additive or super-additive (Figure S5). Thus, linear combinations of Feature and Shape ratings could account for Both-pair ratings for all subjects. Overall, we conclude that information about local features and global shape are integrated linearly (and usually additively) during haptic object perception.

### Local Features and Global Shape are Integrated in a City-Block, and not Euclidean, Manner

Another way of examining how global shape and local features are integrated is to determine which spatial model explains the relationship between the bimodal and unimodal dissimilarity ratings. One possibility is a Euclidean spatial model, where the Both pair ratings would be related to the Shape and Feature ratings by the Pythagorean theorem. The more likely possibility in our dataset is a City-Block spatial model where bimodal distances are an arithmetic sum of their unimodal counterparts (equivalent to the additive model described above; Hyman & Well, 1967). To find the best-fitting spatial model, we calculated the best fitting Minkowski exponent by optimizing the equation:

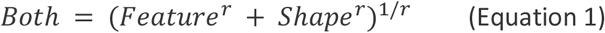

where an r value of 1 would indicate a City-Block, and 2 a Euclidean spatial model.

Figure 3b shows the best-fitting Minkowski exponents for each subject. The median exponent values for C1-C3 were 1.03, 1.14, and 1.20, indicating that the subjects centered around a City-Block-like spatial model. Moreover, the Minkowski exponent was closer to 1 than to 2 for 14 out of 18 subjects (78%). Thus, we conclude that for the majority of subjects, a City-Block spatial model outperformed a Euclidean model in explaining the integration of local features and global shape (Figure 3b).

### Objects that are Rotated Versions of One Another are Perceived as More Similar

C2 included objects that could be perceived as rotated versions of one another. For example, a pair of objects where the shape changes from an “L” shape to inverted-”L” shape could be perceived to be rotated if the local features on index and middle fingers are the same (Figure 4b). If the locations of local features are taken into account while judging changes in global shape, such pairs may be perceived to be more similar than equivalent pairs where different features impinge upon the index and middle fingers. In contrast, if local features are perceived independently from global shape, the same shape change should be rated the same way regardless of the local features (and vice versa for feature changes). To evaluate this possibility, we compared rotated pairs to non-rotated ones. We found, when the Shape change was consistent with a rotation (i.e., going from “L” to inverted-”L”) and the feature change was also consistent with a rotation (in that the same features impinged upon the index and middle fingers), the pair was perceived to be more similar (p < .017; Figure 4b) than when the feature change was incompatible with a rotation (but the shape change was the same). Further, the rotated Both pair was significantly more sub-additive with respect to the sum of the corresponding Feature and Shape ratings compared to non-rotated Both pairs (p < .017; Figure 4a). We did not find a similar effect of rotation for feature changes (p = .343; Figure 4c). Thus, we conclude that objects that were rotations of one another tended to be perceived as more similar than comparable non-rotated pairs.

**Figure 4.**
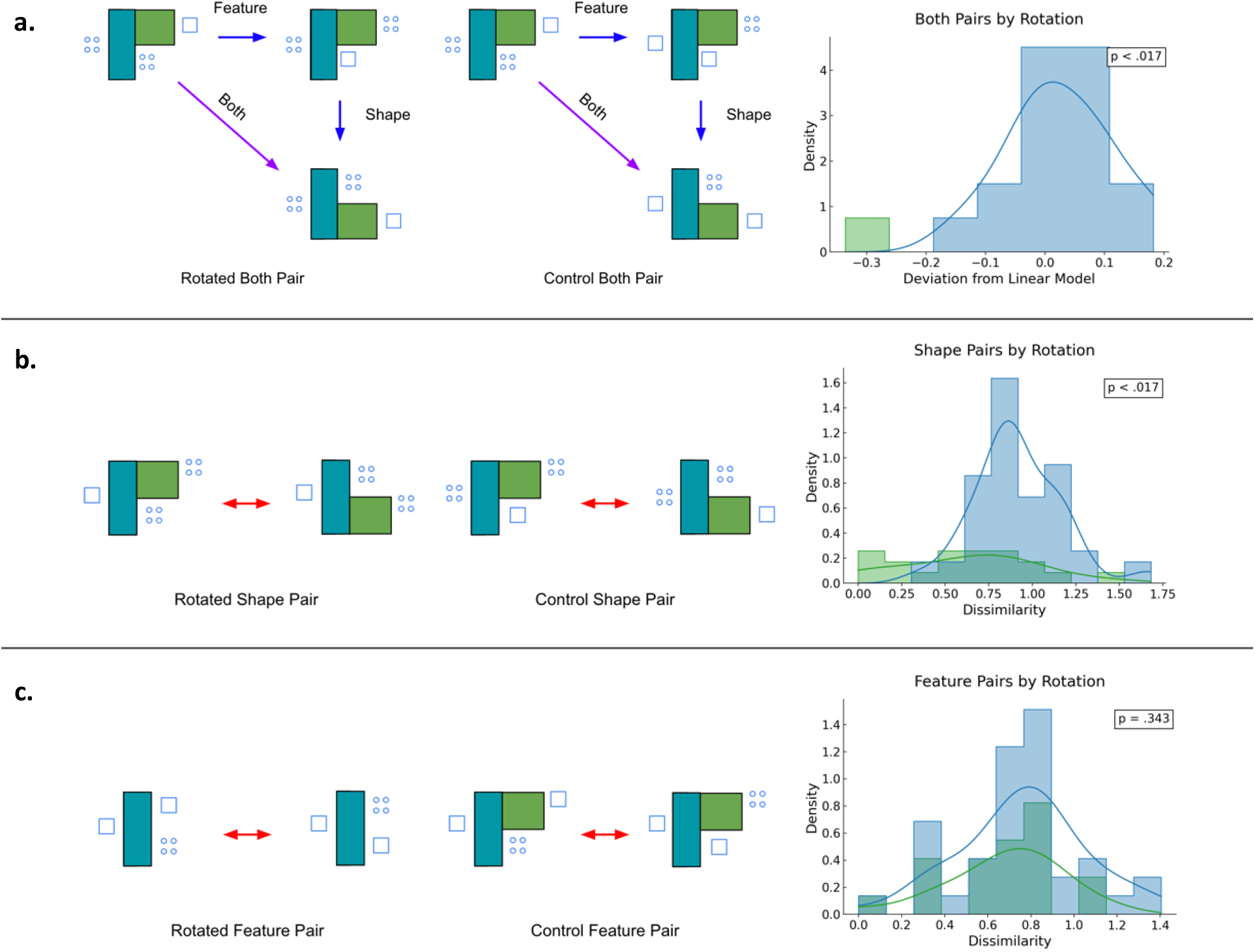
Objects that are rotated versions of one another are perceived to be more similar. a | A sample rotated and control Both pair with corresponding Shape and Feature pairs. When the objects in a Both pair were rotations of each other (versus not), the rating ascribed to the Both pair was sublinear compared to the sum of (non-rotated) Feature and Shape ratings. b | A sample rotated and control Shape pair. Shape changes were rated to be less dissimilar when they were rotations of each other than not. c | Rotated and control Feature pairs. Rotated feature changes were not rated differently from their non-rotated counterparts. The histogram on a shows deviation from a linear model fitted to data averaged across subjects. The histograms in b and c show all ratings by all subjects. p = .05 was chosen as criterion and Bonferroni corrected for multiple comparisons.

## Discussion

Stereognosis requires the integration of tactile signals about local features at each point of contact with proprioceptive signals about the conformation of fingers in space. Here, we aimed to shed light on this integration using a dissimilarity task in a psychophysical experiment. We found that changes in global shape and changes in local features are integrated linearly during haptic object perception. Specifically, for all subjects, perceived dissimilarity when both global shape and local features differed between a pair of objects could be predicted by linear -and for most subjects, additive-combinations of dissimilarity ratings when only global shape or only local features changed. Further, we showed that the City-Block spatial model outperforms the Euclidean spatial model for a majority of the subjects, indicating separable dimensions. These findings enhance our understanding of sensory integration in stereognosis.

Few studies to date have examined the psychophysics of stereognosis. One challenge in studying the psychophysics of stereognosis is disentangling the sensory experience of shape from the effects of semantic encoding. To address this pitfall, Norman et al. (2004) had subjects haptically explore pairs of nonsensical objects (bell-pepper replicas) and indicate whether they were the same shape or different. Nonsensical objects were used to ensure subjects relied only on sensory experiences -rather than semantic categories like “cube”-to make judgments. Subjects could recognize haptically explored objects when these were later presented either haptically or visually, indicating that they were able to achieve an amodal representation of the shape through haptic exploration. Lacey et al. (2007) showed that this ability persists despite changes in viewpoint, suggesting that object shape has a high-order abstract representation in the brain. However, these studies did not systematically investigate the tactile and proprioceptive contributions to stereognosis.

One study that systematically varied texture and object shape in a dissimilarity task similar to ours is by Cooke et al. (2007), who sought to assess the relative contributions of texture and shape to perceived dissimilarity. They found that textures and shape changes were weighted equally when objects were explored haptically, while shape dominated in the visual exploration condition (Cooke et al., 2007). Although this result advances our understanding of the relative weighting of tactile and proprioceptive signals, it does not address the question of which spatial model is appropriate for their integration, as Cooke et al. (2007) conducted their analysis assuming a Euclidean model was appropriate. To our knowledge, ours is the first psychophysical study to directly investigate the spatial model for the integration of proprioceptive and tactile signals.

### Susceptibility to verbal encoding

One possibility is that subjects memorized the ratings ascribed to Shape or Feature changes and reported the sum of the two numbers when both dimensions changed simultaneously. C1 objects were particularly susceptible to this strategy, given that these comprised only three shapes (small, medium, or large spheres) and three features (bumps, holes, or smooth) so both shape and feature could be encoded verbally. However, subsequent experiments comprised more complex objects, making verbal encoding much more challenging. In particular, C3 objects comprised eight shapes and six features (Figure 5) and subjects reported that they did not remember how they rated specific pairs during post-study interviews. Given that the additivity was observed in all object sets, it is unlikely that the observed additivity was an artifact of verbal encoding.

**Figure 5.**
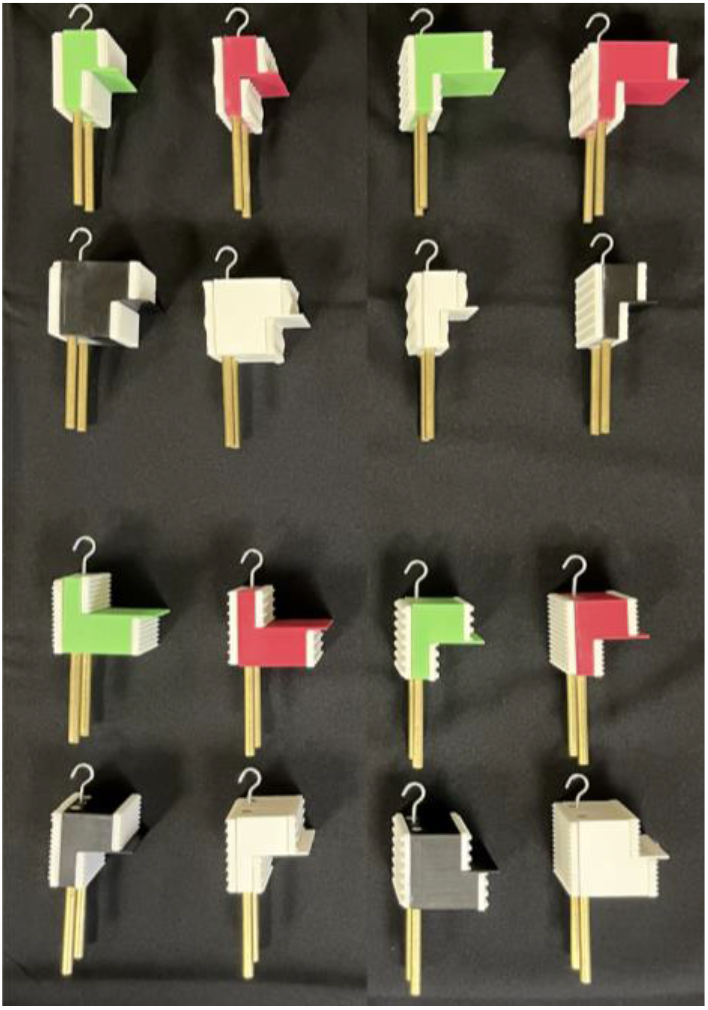
C3 objects. Objects were compared in groups of four, with each group containing a unique feature change (horizontal direction) and a unique shape change (vertical direction).

### Spatial models and separability

We found that the integration of local features and global shape is better described by the City-Block, than the Euclidean, spatial model. It has been argued that two dimensions are integrated in a City-Block manner if they are “separable”, meaning that either one can be attended to while ignoring changes in the other (Garner & Felfoldy, 1970). In contrast, “integral” dimensions which are perceived holistically yield Euclidean distances. For example, in vision, it has been shown that color and shape are separable, while value and chroma are integral dimensions (Hyman & Well, 1967). These psychophysical spatial models are thought to correlate with distances of neural representations (Beeck, Wagemans, Vogels, 2001). Here we showed that unlike value and chroma in vision, tactile and proprioceptive signals are separable dimensions during haptic object perception.

### Object rotations

One of the key questions in neuroscience is how sensory systems achieve stable representations of objects across viewpoints and exploratory procedures. We incidentally observed that rotations of the same object are perceived as being more similar than otherwise equivalent Both pairs. This effect of rotation violates the simple additivity model and implies a higher-level processing of shape, where the object constancy is achieved regardless of the grasp. To perceive two objects whose global shapes are flipped (such as “L” versus inverted-”L” shape) as being rotations of one another, the locations of the local features in space must be interpreted against the change in shape. Indeed, we find that linear combinations of the dissimilarity driven by local features or in global shape alone cannot predict the dissimilarity when both change in a way that can be perceived as a rotation. We only had a limited number of rotated object pairs as this was not the primary focus of our study, but this phenomenon may constitute a fruitful line of future inquiry.

## Methods

### Subjects

A total of 19 subjects participated in the study. Five subjects (three male, two female; mean age = 25.0) participated in C1, six subjects (two male, four female; mean age = 21.8) participated in C2, and eight subjects (two male, six female; mean age = 21.6) participated in C3. All subjects reported normal tactile sensitivity, and none reported any neurological conditions. All experimentation was conducted according to the guidelines set by the Human Institutional Review Board of the University of Chicago. Informed consent was obtained from each subject and every subject was compensated for their participation.

### Objects

All objects were custom designed using Autodesk Fusion 360 software and 3D printed with a FlashForge Creator 3 Pro printer using PLA. For C1, eight sphere shaped objects were printed. The objects had one of three sizes (diameters = 30.0, 50.0, and 70.0 mm) and one of three local features (bumps, holes, or smooth surface; Figure S1A). We included all size-feature combinations except for Large-holes. For the local features, “bumps” were 2.0 mm tall cones with round tips (base diameter = 6.0 mm) which tiled the sphere surface with a center-to-center distance of ∼9 mm. Holes were semi-spherical indentations (diameter = 8.0 mm) that were distributed with a center-to-center distance of ∼9 mm.

C2 objects were 12 square-based blocks of different compositions. These objects were designed such that the thumb would contact one side (the square base), and the index and middle fingers would contact the surface on the opposite side, where a ledge separated the regions for the index and middle fingers (Figure S1b). Three shapes were possible: both index and middle fingers 18.0 mm away from the thumb (“I” shape), index 39.5 mm away and middle 18.0 mm away from the thumb (inverted-”L” shape), and index 18.0 mm away and middle 39.5 mm away from the thumb (“L” shape). The base (i.e., the thumb side) was 50.8 × 46.5 mm. There were two possible local features (bumps and smooth surface), and these were distributed across the three surfaces to yield different combinations. We included four feature combinations (in order of thumb, index, and middle: bump-bump-smooth, bump-smooth-bump, smooth-bump-bump, and smooth-smooth-bump). Bumps were half-sphere protrusions with a diameter of 5.0 mm and tiled the surface with a center-to-center distance of 7.1 mm.

C3 objects were shaped similarly to C2, but the spacing between the fingers was varied. We constructed eight shapes by varying the thumb-index and thumb-middle distances to different degrees within the range of 22.0 - 54.0 mm (Figure S1c; Figure 5). The base surface that the thumb contacted was 46.0 mm long and 50.0 mm tall. There were six different local features (*Small dots*: 1.5 mm radius, 7.0 mm center-to-center; *Horizontal/vertical bars*: 1.5 mm tall, 1 mm wide, 6 mm center-to-center; *Square dots*: 1.5 mm tall, 1.0 mm wide, 3.25 mm center-to-center; *Waves*: 3.0 mm peak-trough distance, 1.78 peaks per cm^2^ area; *Cones*: 2.5 mm high, 5.0 mm base diameter, 2.25 mm top diameter, 6.5 mm center-to-center). These features were printed separately with a Formlabs Form 3+ resin printer using white resin and attached to the objects with tape. Each object had one type of local feature.

### Dissimilarity task

All object sets were tested using the same dissimilarity task. Subjects were seated at an experimental table where they could not see the objects (in C1 this was accomplished by instructing the subjects to keep their eyes closed; a curtain was used in C2 - 3). On each trial, subjects were tasked with grasping a pair of objects in sequence and giving a rating of dissimilarity. For C1, objects were held slightly above the open right palm of the subject, and after a cue the subject grasped the object with their thumb, index, and middle fingertips by flexing their fingers. Subjects were instructed to not move their fingers around the object and to continue grasping until a second cue. For C2 - 3, objects were mounted vertically on a disc on the experimental table, and subjects reached forward and grasped the objects with their right hands (Figure 1b). The thumb contacted the left and the index and middle fingers contacted the right side of the object. In all experiments, subjects grasped each object for approximately three seconds, and the time between the presentation of two objects in a pair was approximately 2-4 seconds.

Minimal instructions were provided on how to give ratings. All subjects were asked to rate how dissimilar the pair of objects were “overall”, and to give a rating of zero if the objects were the same. Subjects were instructed to “rate a pair of objects that is twice as dissimilar as another pair twice as high as the other pair”. Subjects were free to use their own scales.

C1 had eight different objects, and these were parametrically combined to yield 36 total pairs (eight “Same” pairs, seven “Feature” pairs, seven “Shape” pairs, and 14 “Both” pairs). Data were collected in five sets, each consisting of 36 pairs (180 trials total). Sets were interleaved with short breaks. C2 had 12 objects that were combined to form 78 pairs (12 Same, 18 Feature, 12 Shape, and 36 Both pairs). Two subjects completed five repetitions (390 trials), and the remaining four completed three repetitions (234 trials). C3 had 16 objects, which were paired with one another in groups of four objects that contained a specific feature change, shape change, and both change comparison. This yielded 42 pairs (six Same, 12

Feature, eight Shape, eight Both, and eight Catch pairs). Two subjects completed four repetitions (168 trials) and the remaining six completed three repetitions (126 trials). Same trials were included as controls. In C3, some trials also included “catch” objects which were designed to be different from the experimental objects. These were included as distractors.

### Data Analysis

Because subjects were instructed to give a rating of zero if the objects were the same, for subjects who never gave a rating of zero, the lowest rating was subtracted from all other ratings. This was the case for three out of 19 subjects (one of which was later excluded). All ratings were normalized by dividing by the mean rating of the set (excluding Same or Catch pairs). For each pair, a dissimilarity score was calculated by averaging the normalized dissimilarity ratings across sets.

For each subject, we computed a signal-to-noise (SNR) value by dividing the standard deviation of mean dissimilarity ratings across all pairs (except for Same and Catch pairs) by the average standard deviation in dissimilarity ratings given to each pair. To the extent that subjects varied their ratings across pairs more than within pairs, the SNR value was greater than 1.

We fitted a linear regression model with a forced zero intercept to predict dissimilarity scores for when both local features and global shape changed based on the sum of the dissimilarity scores for corresponding Feature and Shape pairs. The regression model yielded two measures: the coefficient of determination, which helped us determine whether the Both ratings were predictable from a linear sum of the Shape and Feature ratings; and the slope, which determined whether the relationship was additive. Because the R^2^ value was affected by the spread of the data while we were primarily interested in whether the Both ratings equaled the summed values, we devised a second error metric by dividing the deviation from the fitted model (root mean squared error) by the intrinsic noise in the data. We quantified the noise as the RMS of the deviation of individual ratings from the mean rating of the corresponding pair (excluding Same and Catch pairs). For each subject and pair, we computed the RMS deviation of individual ratings from the mean rating of that pair. We then averaged the RMS deviation values across pairs to get a noise estimate.

To estimate the best-fitting Minkowski exponent, we iteratively optimized Equation 1 between r values of 0.1 and 10. We used the L-BFGS-B optimization algorithm with the “minimize” function from the SciPy package for Python with a mean squared error loss function (Virtanen et al., 2020).

Finally, for each subject, performed Monte Carlo simulations under the assumption that Feature and Shape ratings are combined additively (Figure S4) or linearly (Figure S5) to predict Both-pair ratings. For this, we plotted the sum of Feature and Shape pair ratings against themselves (factored by a coefficient for the linear model), and then injected randomly sampled noise to both axes. The noise was sampled from a distribution of deviations of each individual rating from the mean rating of the corresponding pair (yielding n x p values where n is the number of blocks and p the number of pairs). We sampled three values from this distribution, averaged them, and added this to each data point. We injected noise separately to Shape ratings, Feature ratings, and the sum of the Shape and Feature ratings. We then summed the noisy Shape and Feature ratings to simulate a distribution of summed values, and plotted these against the noisy sum of Shape and Feature ratings (which simulated Both pair ratings). For each simulated model, we computed the error of an additive or linear model (Deviation / Noise). For the linear model we used the same coefficient as the best fitting linear model of the original ratings. We repeated this 10,000 times per subject, and compared the empirical error of the relevant model to the distribution of simulated error values. p < .01 was used as criterion.

## Supplementary Figures

**Figure S1.**
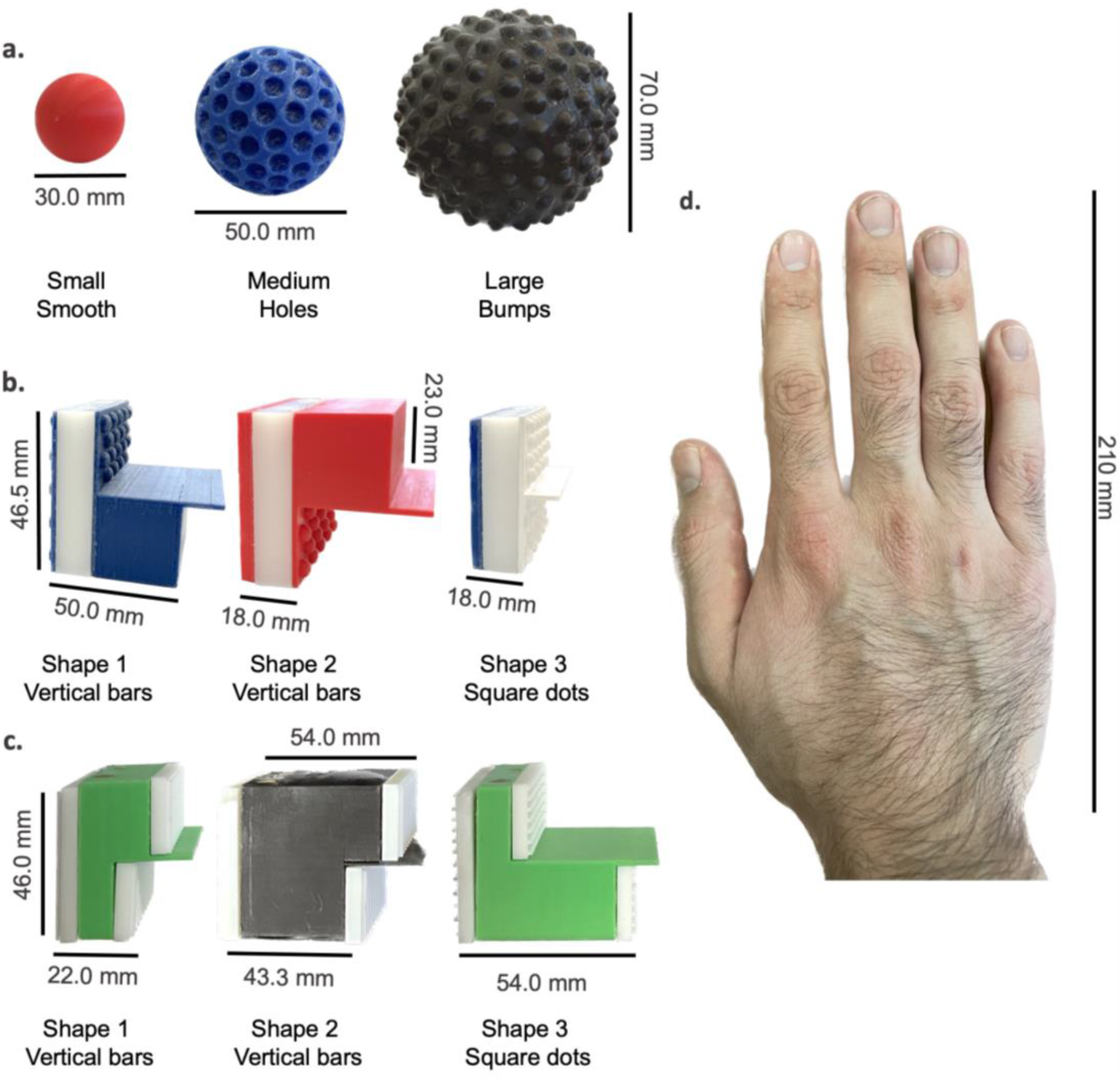
Objects. a | Sample C1 objects showing the three sizes and three local features. These size and local features were combined to create eight different objects. b | Sample C2 objects, showing the three global shapes and three of the local feature patterns. There was a total of four unique local feature patterns (“smooth-smooth-bump” not shown), and these were combined with the three shapes to create 12 objects. c | Sample C3 objects. There was a total of eight global shapes and six local features, which were locally combined to create 16 objects. d | Human hand for scale.

**Figure S2.**
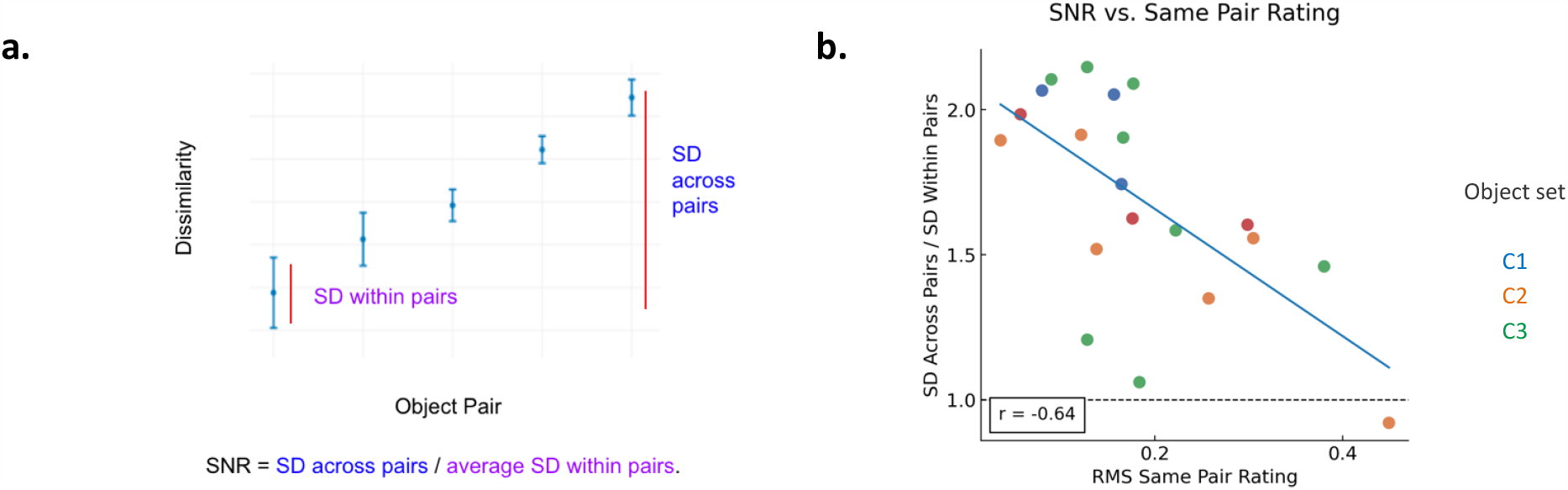
Signal-to-noise ratio negatively correlates with deviation from zero of Same pair ratings. a | Graphical representation of the signal to noise (SNR) metric. SNR was defined as standard deviation of mean ratings across all pairs divided by the average standard deviation of ratings within pairs. b | SNR vs root mean squared Same pair rating by subject and object set. For C1, two of the subjects which were not presented Same pairs are not shown. Both of these subjects also had SNR values greater than one.

**Figure S3.**
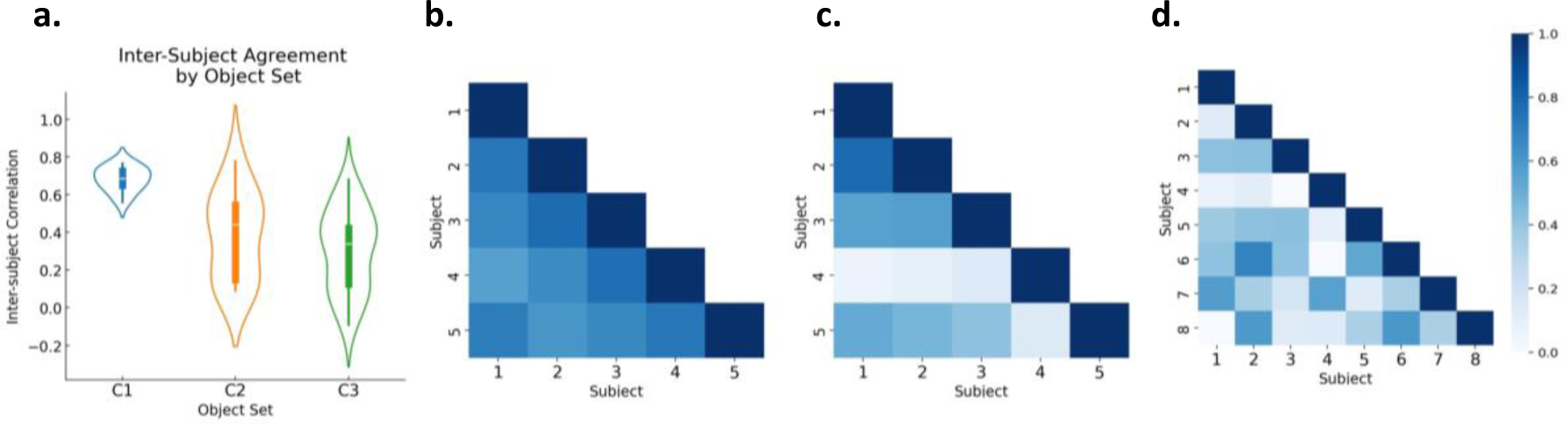
Inter-subject agreement across object sets. a | Violin plot showing the inter-subject correlation coefficients by object set. The mean correlation coefficients are 0.68, 0.38, and 0.31, for C1-C3, respectively, highlighting the increasing complexity of the objects from C1 to C3. b, c, d | Correlation matrices for C1, C2 and C3, respectively. Note that dissimilarity ratings were normalized to the average of the respective category (Feature, Shape, or Both) and Same pairs were excluded to prevent the subjects’ combination strategies (e.g., additive or subadditive) affecting the correlation values.

**Figure S4.**
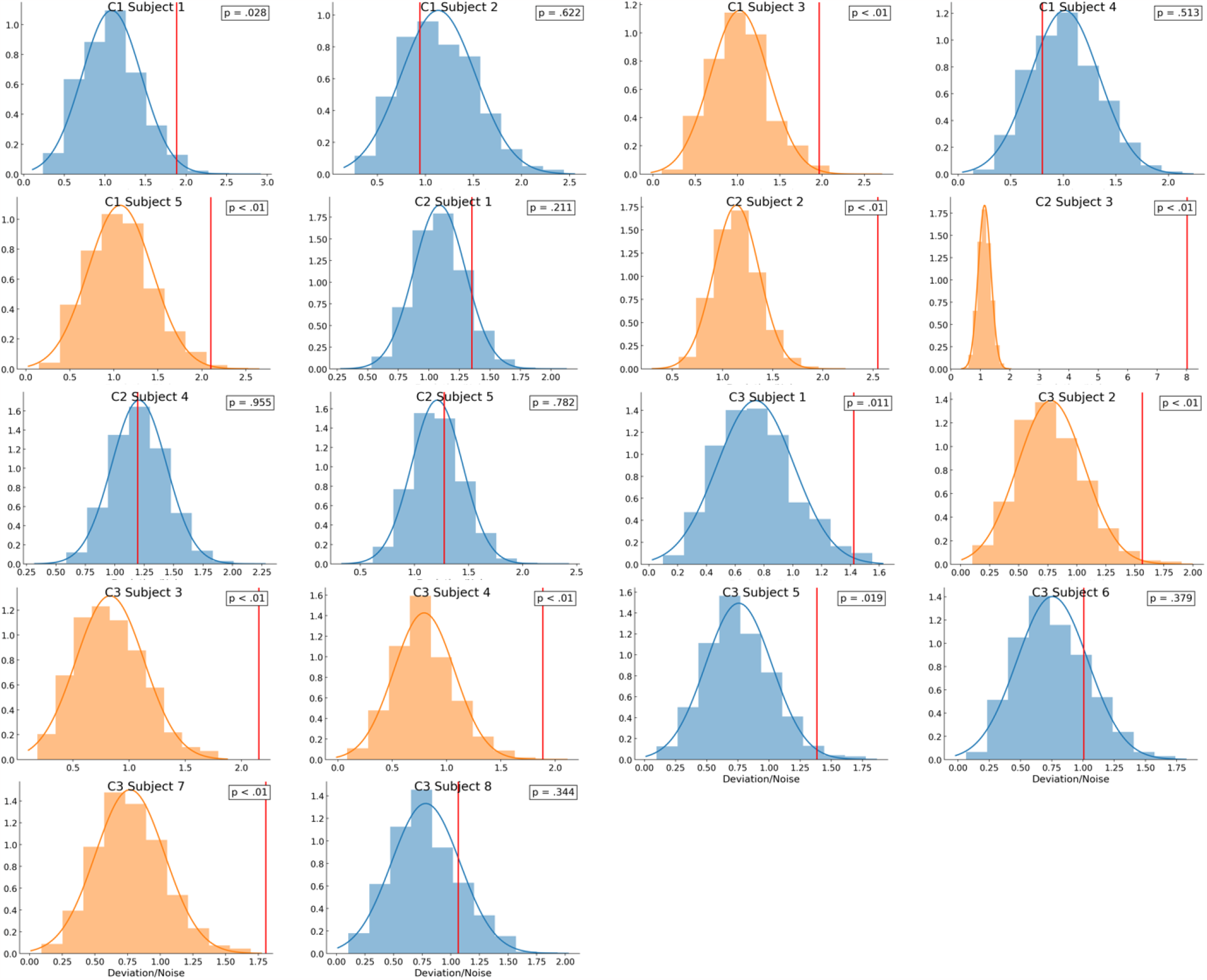
Monte Carlo simulated error distributions assuming additivity versus actual error for individual subjects in C1-3. For each subject, a simulated additive model was created by plotting the sum of Feature and Shape pair ratings against themselves, and then injecting randomly sampled noise to both axes. The noise was sampled from the set of deviations of each rating from the mean rating of that pair across blocks. For each simulated model, we computed the error of the additive model (Deviation / Noise). We repeated this 10,000 times per subject, and compared the empirical error of the additive model to the distribution of simulated error values. p < .01 was used as criterion. Subjects with significant deviations from additivity are shown in orange.

**Figure S5.**
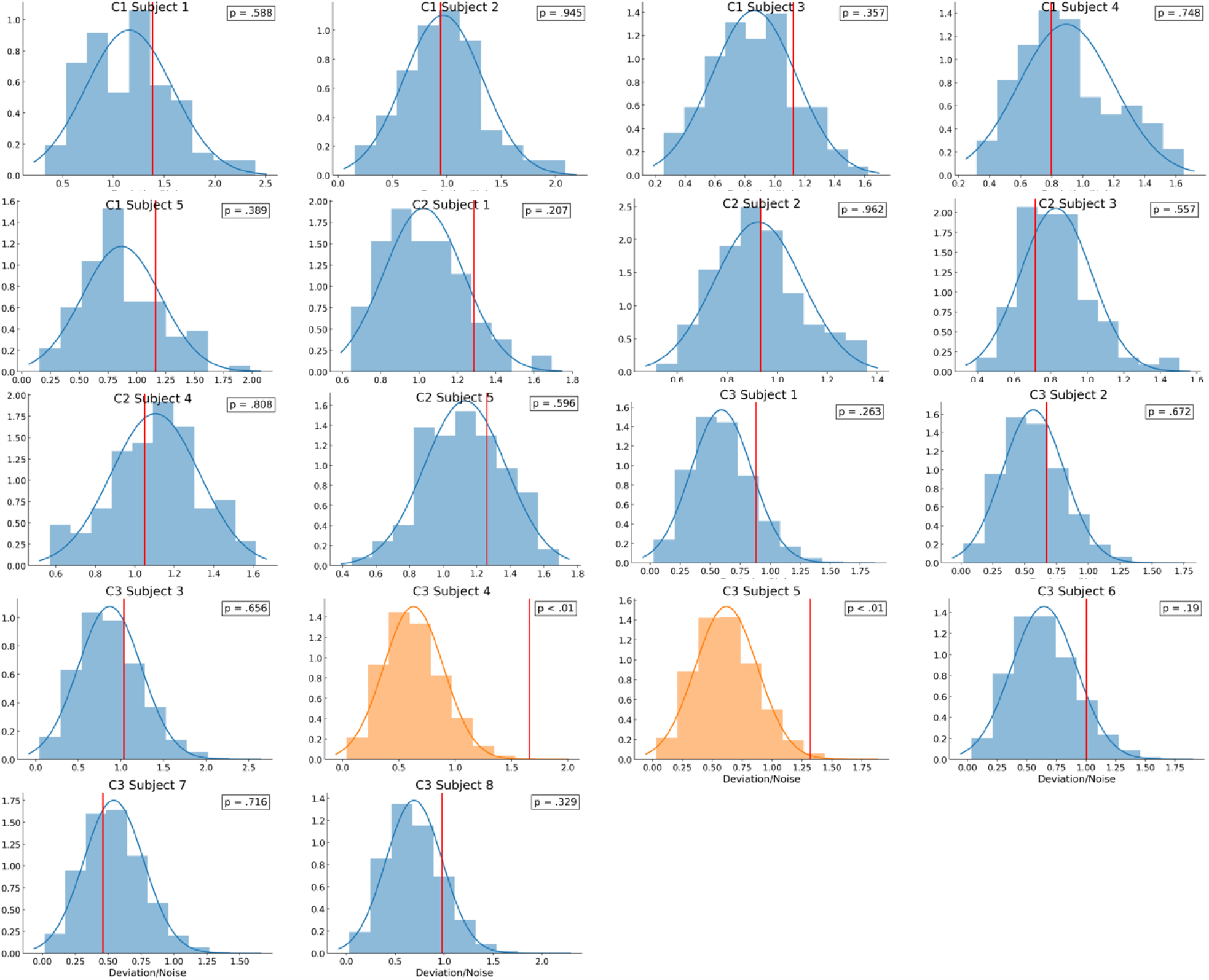
Error of the best fitting linear model compared to Monte Carlo simulated error distributions with the same model. For each subject, we first fitted a linear regression model through the intercept to predict Both pair ratings from the sum of Shape and Feature ratings, which yielded a best-fitting coefficient for the summed values. We then simulated a linear model taking this coefficient to be ground truth. The rest of the process was the same as in Figure S4. Only two of the subjects significantly deviated from linearity (p > .01).

**Figure S6.**
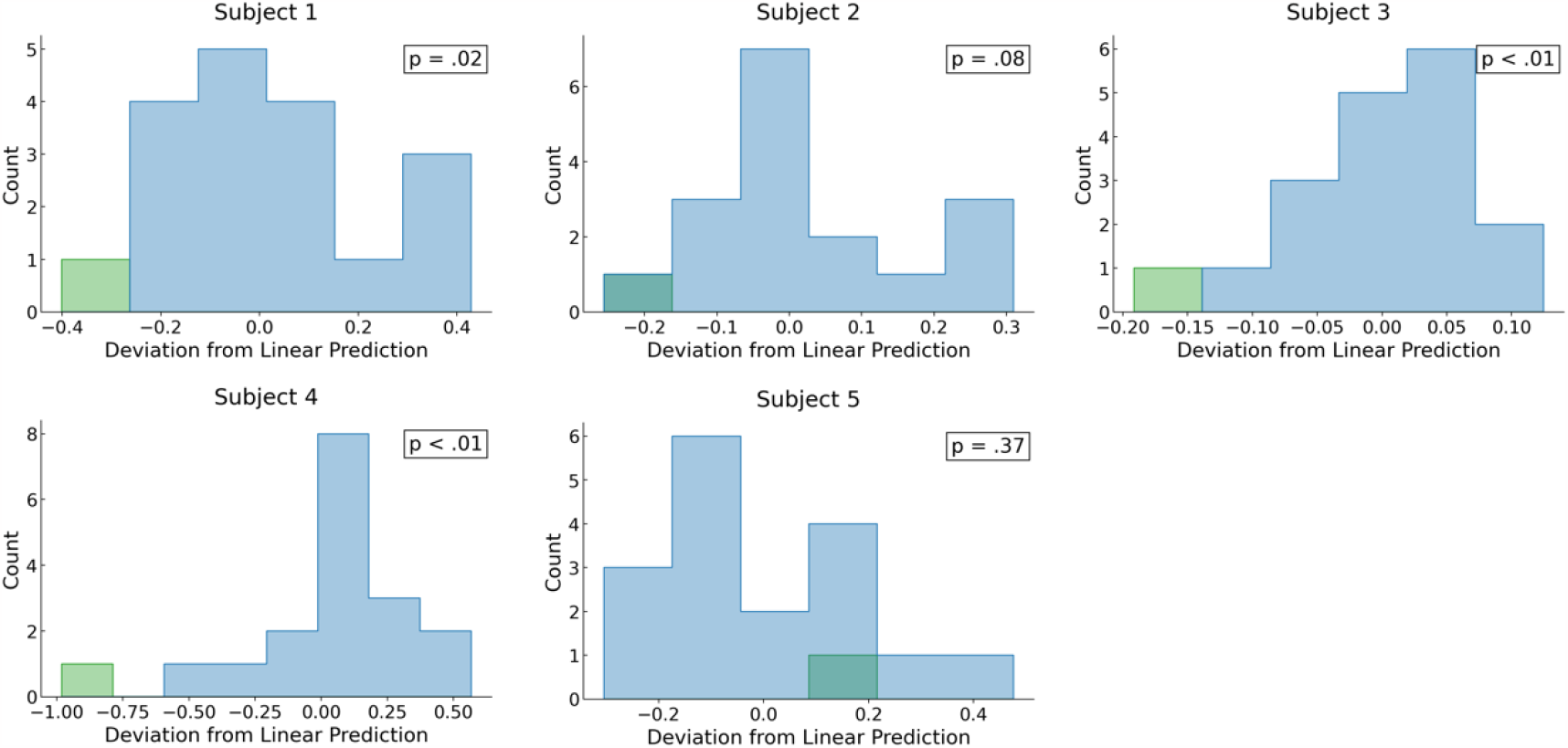
Both pairs where the objects are rotations of each other are perceived as less dissimilar. Both pairs that were rotations were significantly more subadditive compared to non-rotated Both pairs for two out of five subjects in C2. p < .01 was used as criterion.

